# PP2A inhibition instructs spliceosome phosphorylation to create splicing vulnerability in colon adenocarcinoma

**DOI:** 10.1101/2023.07.12.548685

**Authors:** Matheus H. Dias, Vladyslava Liudkovska, Onno B. Bleijerveld, Arno Velds, René Bernards, Maciej Cieśla

## Abstract

Protein phosphatase 2A (PP2A) is a versatile enzyme affecting many aspects of cellular physiology. However, it remains unclear which of the cellular processes are most perturbed upon PP2A inhibition and how these perturbations could be exploited therapeutically. We report an unanticipated sensitivity of the splicing machinery to phosphorylation changes in response to PP2A inhibition by LB-100 in colorectal adenocarcinoma. We observe enrichment for differentially phosphorylated sites within cancer-critical splicing nodes of U2 snRNP, SRSF and hnRNP proteins. Altered phosphorylation endows LB-100-treated colorectal adenocarcinoma cells with differential splicing patterns. In PP2A-inhibited cells, over 1000 events of exon skipping and intron retention affect regulators of genomic integrity. Finally, LB-100-evoked alternative splicing is predicted to be a source of neoantigens that can improve cancer treatment responses to immune modulators. Our findings provide a potential explanation for the pre-clinical and clinical observations that PP2A inhibition sensitizes cancer cells to immune checkpoint blockade and genotoxic agents.

## INTRODUCTION

Protein phosphatase 2A (PP2A) is a pleiotropic serine/threonine phosphatase that acts in a multitude of cellular signaling pathways^1^. PP2A is considered a tumor suppressor gene given their potential to modulate various cancer-related signaling pathways and thereby inhibit tumor development^2, 3^. Drug development efforts have therefore focused on small molecule activators of PP2A, as such drugs could potentially act on multiple cancer-relevant signaling routes to suppress tumorigenicity. Surprisingly, inhibition of PP2A with the small molecule inhibitor LB-100 has also exhibited anti-cancer effects, especially when used in combination with radiotherapy or specific chemotherapy drugs^4^. This effect is believed to result from the notion that PP2A inhibition affects DNA repair mechanisms, making cells more vulnerable to DNA damaging agents^5, 6^. Our recent data show increased oncogenic signaling in response to PP2A inhibition, causing DNA replication stress, which may also contribute to a greater sensitivity to DNA-damaging agents^7^. An independent and potentially powerful anti-cancer effect of PP2A inhibition stems from the finding that LB-100 also enhances the efficacy of immune checkpoint blockade (ICB) in different cancer models by fostering T-cell and cGAS-STING activation^8, 9^. Moreover, PP2A inhibition has been shown to turn immunologically “cold” tumors (i. e. those poorly recognized by immune system) “hot” by inhibiting DNA mismatch repair^10^. Such “promiscuous” sensitization of PP2A inhibition to different therapeutic modalities reflects the central role of this phosphatase in controlling cancer cell homeostasis.

One of the emerging therapeutic strategies for cancer treatment is to leverage tumor-associated epitopes presented in the context of Major Histocompatibility Complex Class I (MHC-I) ^11, 12^. These non-canonical immunopeptides can activate the immune system against cancer cells through recognition by autologous T cells. The newly formed neoepitopes, which are not immune-privileged and are expressed exclusively outside the healthy tissues, create an attractive window for anti-cancer vaccines or checkpoint immunotherapies^11^. The cancer-specific formation of neoantigens has been reported to stem from somatic mutations and genetic alterations^11^, translation from novel or non-canonical open reading frames^13^, and altered function of splicing components^14, 15^. Regarding the latter, various cancers have been found to have mutations in the core splicing machinery^16^. It is now acknowledged that alternative splicing can be a source of tumor neoantigens, and there are ongoing efforts to develop pharmacological treatments that target this process^17, 18^. Accordingly, a recent study indicates that pharmacologic modulation of splicing drives the formation of functional neoantigens to elicit anti-tumor immunity, inhibiting tumor growth and enhancing checkpoint blockade in a manner dependent on host T cells^14^. Collectively, these data suggest that splicing modulators are attractive targets for combinatorial therapies alongside immune checkpoint blockade approaches.

Given the growing body of evidence supporting the anti-cancer effects of LB-100, particularly its capability to enhance tumor immune sensitization^8^, understanding the global impact of this PP2A inhibitor on the phosphoproteome of cancer cells could provide further mechanistic insights into why cancer cells are susceptible to LB-100. Moreover, such studies could suggest potent drug combinations by identifying specific processes that are perturbed by the drug. Here, we addressed the impact of LB-100 on the phosphoproteome of colorectal cancer cells to infer the most relevant cellular processes altered by the drug. Our results identify a major role for PP2A in the proper execution of RNA splicing. As alternative RNA splicing can yield neoantigens on cancer cells, our data provide a further rationale for combining PP2A inhibition with checkpoint immunotherapies.

## RESULTS

### PP2A inhibition drives changes in the phosphoproteome of splicing regulators

To determine proteome dynamics upon acute inhibition of PP2A in cancer cells, we first treated human colorectal adenocarcinoma cell line SW-480 with LB-100 for 12 hours (Figure 1A). We subsequently performed paralleled quantitative mass spectrometry-based proteomics and phosphoproteomics to delineate how the protein repertoire is affected by PP2A inhibition. Comparative analysis of LB-100-treated samples with controls revealed relatively few changes in protein abundance, with 120 and 239 proteins being respectively down and up-regulated in response to PP2A inhibition (Figure 1B). This result is in stark contrast with phosphoproteomic analysis that revealed widespread changes in protein phosphorylation status, with >1700 proteins undergoing differential phosphorylation in LB-100-treated cells (Figure 1C). Consistently with the expectation that inhibition of phosphatase activity would predominantly evoke hyperphosphorylation changes, we observed bias toward this type of phosphorylation events (62%, encompassing 1069 significantly differentially phosphorylated proteins with 2099 significantly affected phosphosites), as compared to hypophosphorylation events (38%, comprising 649 proteins with 1364 affected phosphosites) in LB-100-treated cells.

**Figure 1.**
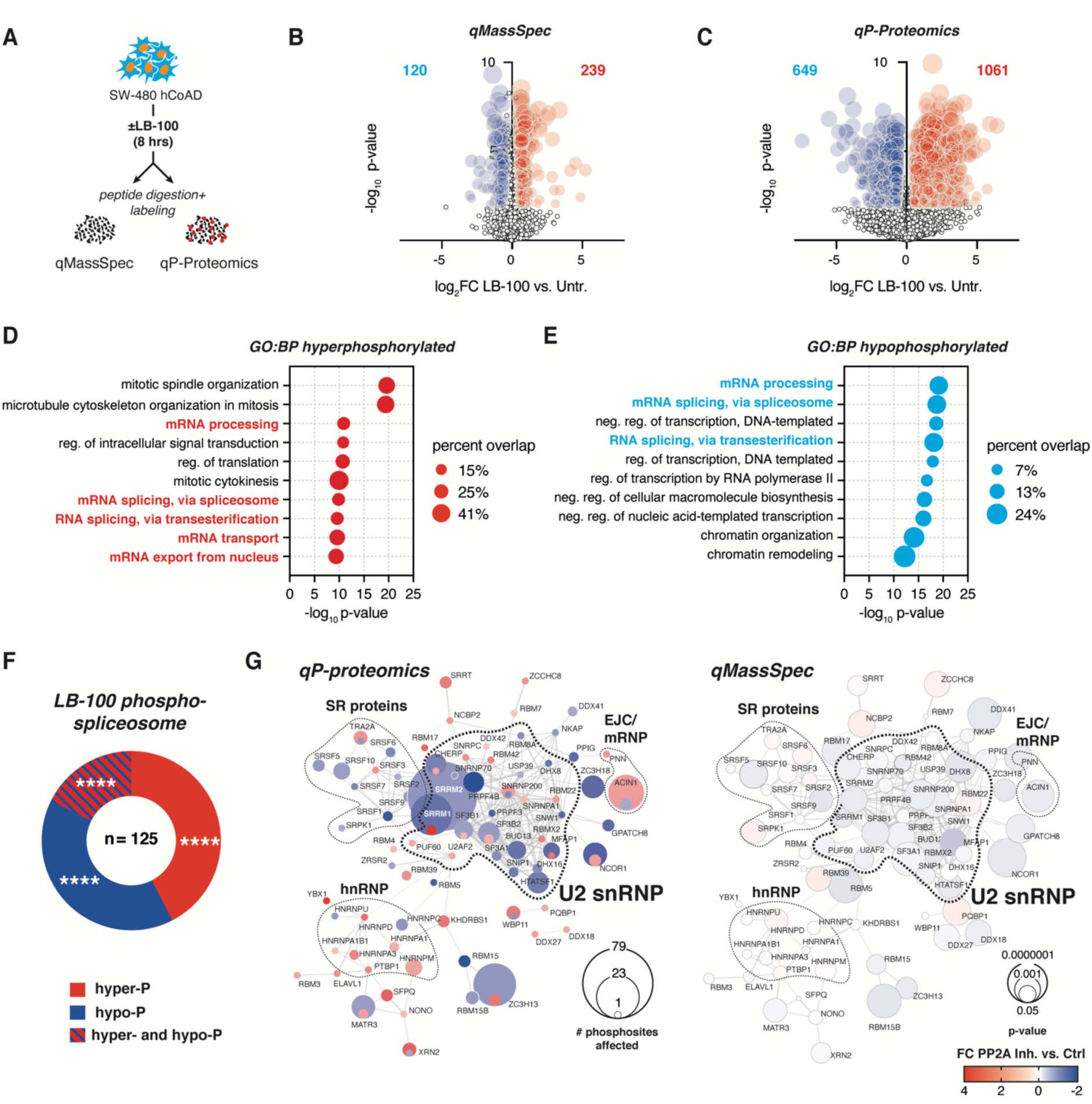
PP2A small drug inhibition by LB-100 drives phosphoproteome changes enriched for multiple components of spliceosome. **(A)** Schematic illustrates experimental approach to determine proteomic (quantitative Mass Spectrometry, qMassSpec) and phosphoproteomic (qP-Proteomics) changes in response to PP2A inhibitor LB-100 in human colorectal adenocarcinoma (hCoAD) cell line SW-480. **(B)** Volcano plot shows protein expression changes in response to 4μM LB-100 stimulation (8 hrs) compared to untreated controls (Untr.). Upregulated (red) and downregulated (blue) proteins with False Discovery Rate (FDR)<0.05 and |Log_2_FoldChange|>0.58 are shown. **(C)** Volcano plot shows phosphorylation changes in response to 4μM LB-100 stimulation (8 hrs) compared to untreated controls. Hyper-(red) and hypo-phosphorylated (blue) events with FDR<0.05 and |Log_2_FC|>0.58 are shown. **(D)** Gene Ontology (GO) analysis for 1061 hyper-phosphorylated proteins. BP – Biological Process. **(E)** GO analysis for 649 hypo-phosphorylated proteins. **(F)** Donut chart shows distribution of proteins with significantly changed (FDR<0.05 and |Log_2_FC|>0.58) phosphorylation status. ****p<0.0001 (hypergeometric test vs. background). **(G)** Interaction maps of spliceosome components hyper- and hypo-phosphorylated (left) in response to 8 hrs of LB-100 stimulation in SW-480 cells. Corresponding analysis of protein expression changes is shown (right). Connecting lines show interactions from STRINGdb. See also Figure S1 and Tables S1 and S2.

We reasoned that changes in phosphorylation downstream to PP2A inhibition may affect disparate protein families important for cancer cell viability. Indeed, analysis of proteins exhibiting increased (Figure 1D) and decreased (Figure 1E) phosphorylation upon LB-100 treatment revealed enrichment for pathways critical to cancer, including DNA replication, cell cycle progression, signal transduction, and chromatin remodeling (Figures 1D and E). Strikingly, among the most overrepresented protein families in both hyper- and hypophosphorylated groups, we detected numerous factors linked to RNA metabolism, with an unanticipated enrichment for proteins related to mRNA splicing regulation and control (Figures 1D-F). Indeed, approximately 39% (125 out of 315) of known human splicing regulators were differentially phosphorylated in response to LB-100 treatment. We observed a significant enrichment for all categories of phosphorylation events (hyper-, hypo-, and biphasic phosphorylation encompassing splicing regulators in which different amino acid residues underwent increased or decreased phosphorylation), rather than skewing for hyperphosphorylation events observed for the other groups of aberrantly phosphorylated proteins (Figure 1F).

The spliceosome is a megadalton, multimodal RNA-protein machinery consisting of 5 small nuclear ribonucleoprotein complexes and a multitude of transiently interacting splicing regulators that can both activate and repress alternative splicing^19^. Projection of differentially phosphorylated splicing factors on a map of protein-protein interactions within the spliceosome unveiled specific groups of splicing regulators sensitive to phosphorylation changes induced by PP2A inhibition (Figures 1G and S1A-D). These phosphorylation changes were largely uncoupled from differences in total protein abundance, in line with previous reports that protein phosphorylation affects function and protein-protein interactions rather than expression of the differentially phosphorylated protein. We observed that multiple constituents of the core complex of U2 small nuclear ribonucleoprotein (snRNP) underwent differential phosphorylation events in response to LB-100. U2 snRNP is recruited to the intron branch site to form the A complex and then, upon recruitment of U4/U6.U5 tri-snRNP, it forms the B and B^act^ splicing complexes^19^. In this way, U2 snRNP binding confers a rate-limiting and fidelity-ensuring stage of the reaction that enables nucleophilic attack of a branch point adenosine on the 5’ splice site. We observed that U2 components previously described as key cancer fate determinants^18, 20^, such as RBM8A, splicing factor 3B subunit 1 (SF3B1) or U2AF1 were among mis-phosphorylated proteins (Figure 1G). Moreover, we found that many of the differentially phosphorylated proteins belong to the serine-arginine-rich splicing factor family (SRSF) and heterogenous nuclear RNPs (hnRNPs) (Figures 1G and S1B). SRSFs and hnRNPs are generally recognized as, respectively, activators and repressors of the splicing and key determinants of cancer progression through concerted control of cancer-initiating alternative splicing events^21^. Proteins such as SRSF1 or SRSF2, which we observe to be mis-phosphorylated in PP2A-inhibited cells are known cancer regulators^21^. Of note, for the four splicing proteins found to be recurrently mutated in cancer (SRSF2, U2AF1, SF3B1, and ZRSR2), we observed changes in the phosphorylation patterns in response to LB-100. Mutations often result in abnormal activity of these splicing factors and altered splicing patterns^18^, contributing to the oncogenic phenotype. However, how the phosphorylation of these factors might change their function during cancer progression and in response to targeted therapies remains largely unexplored.

### PP2A inhibition promotes alternative splicing in colorectal adenocarcinoma cells

Motivated by the observations that phosphorylation critically tunes protein-protein interactions and affects protein activity, we reasoned that the observed rewiring of splicing factor phosphorylation might affect splicing outcomes in colorectal adenocarcinoma cells treated with LB-100. To address this, we performed paired-end RNA sequencing (RNA-seq) in control and LB-100-treated cells at the time points matching our prior quantitative phosphoproteomic analysis. To rule out possible cell-specific effects, we performed sequencing in two different human colorectal adenocarcinoma cell lines having different genetic backgrounds (KRAS mutation for SW-480; BRAF mutation for HT-29, Figure 2A). Analysis of independent duplicate RNA-seq experiments using the rMATS pipeline^22^ identified reproducible alterations within ∼2,000 alternative splicing events (ASEs) enriched predominantly for exon skipping (SE) and intron retention (IR) (Figure 2B). Using delta percent spliced-in (ΔPSI) as a metric for splicing efficiency, we observed that LB-100 treatment led to a significantly higher number of ASE. That resulted in increased inclusion of alternative exons and aberrant intron retention as compared to control cells (Figures 2B and C). This entails that PP2A inhibition favors alternative splicing and impacts ASE patterns by abnormal functionality of core splicing machinery. This could be achieved either by boosting the function of splicing activators that recognize specific regulatory motifs in alternative exons with increased splicing efficiencies, or by decreasing the activity of negative regulators. This is in line with our observations on the enrichment of SRSF splicing activators among hypophosphorylated proteins in response to LB-100 (Figures 1G and S1B).

**Figure 2.**
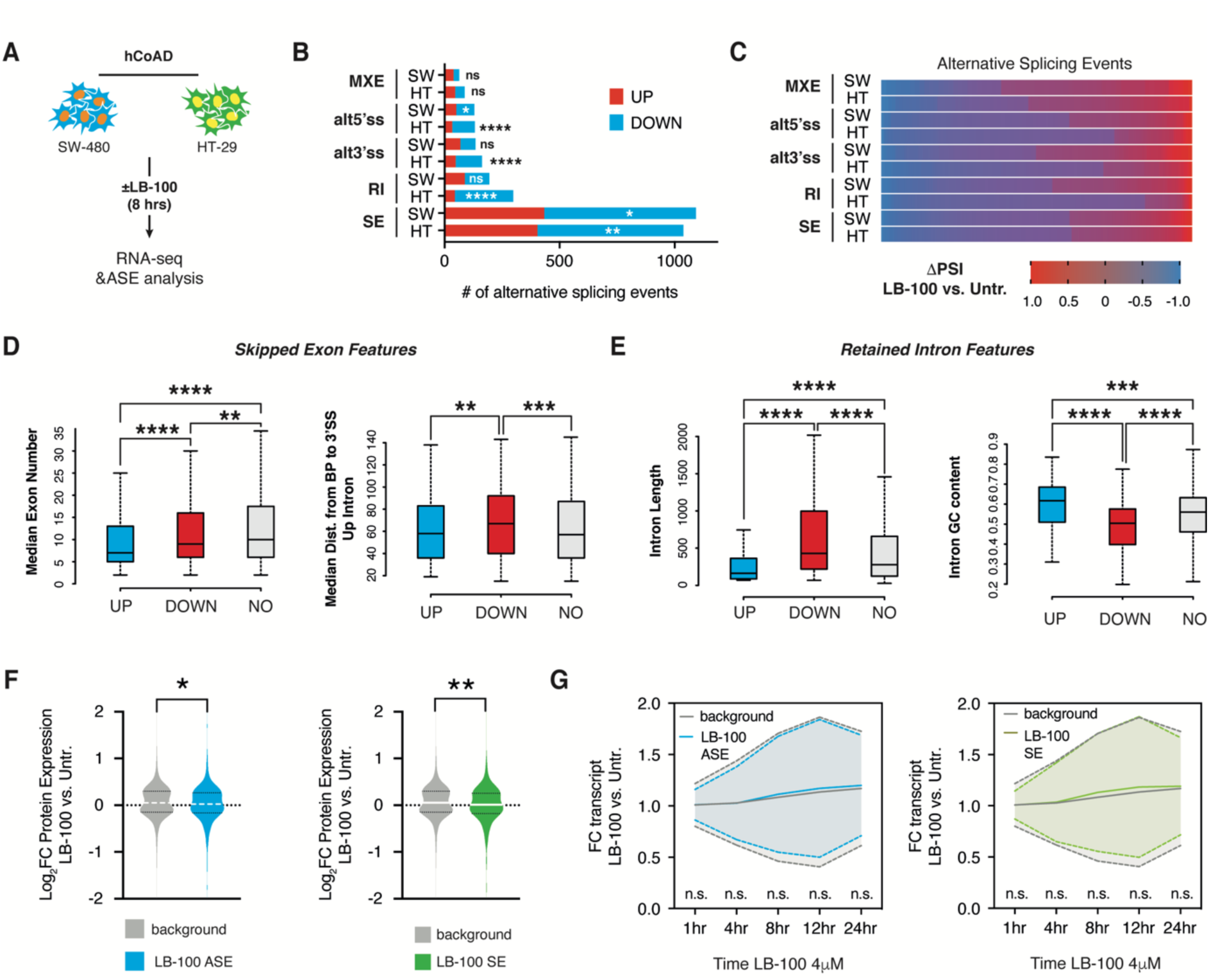
Selective inhibition of PP2A impacts alternative splicing patterns in colorectal adenocarcinoma. **(A)** Experimental setup employed to capture splicing changes in response to PP2A inhibitor LB-100 in human colorectal adenocarcinoma (hCoAD) cell lines SW-480 and HT-29. **(B)** Bar graph shows numbers of skipped exons (SE), mutually exclusive exons (MXE), alternative 3’ splice sites (alt3SS), alternative 5’ splice sites (alt5SS), and retained introns (RI) in LB-100-treated SW-480 and HT-29 cells. ****p<0.0001; ***p<0.001; **p<0.01 in up- vs. down-spliced mRNAs in each family (hypergeometric test). **(C)** Heatmap illustrates quantitative extent of delta percent spliced-in (ΔPSI) for alternative splicing events significantly mis-spliced in response to LB-100 stimulation (FDR<0.05 and |ΔPSI|>0.1) **(D)** Box-and-whisker plots showing significant differences in median exon number and distance of Branch Point (BP) ^35^ to 3’splice site (3’ss) in downstream intron in exons with decreased (n=641) or increased (n=409) percent spliced-in values. ****p<0.0001; ***p<0.001; **p<0.01 (Mann-Whitney U test). **(E)** Box-and-whisker plots showing significant differences in median intron length and guanine-cytosine (GC) content in introns with decreased (n=105) or increased (n=87) percent spliced-in values. ****p<0.0001; ***p<0.001 (Mann-Whitney U test). **(F)** Violin plot shows decreased levels of proteins resulting from all alternatively spliced transcripts (left) and in exon skipping group of ASE (right) in 4 μM LB-100-treated SW-480 cells compared to control. **p<0.01 (t test). **(C)** Graphs show time-resolved changes in all alternatively spliced transcripts (left) and mRNAs from skipped exon group (right) in response to LB-100 stimulation. Mean expression is shown as a solid line ± SD depicted as dotted lines **p<0.01; *p<0.05 (one-way ANOVA). See also Figure S2 and Tables S3 and S4.

Specific regulatory motifs within alternatively spliced mRNAs may be determinants of splicing outcomes in response to the changed activity of spliceosome. In line with that notion, differentially spliced exons in LB-100-treated cells shared specific features contributing to alternative exon usage such as their short length or increased distance from branch point adenosine to the 3’splice site (Figure 2D). Instead, for introns with increased retention scores in response to LB-100 treatment, we observed a significant tendency for increased length and lower GC content, features classically associated with the difficult-to-recognize intervening sequences (Figure 2E). Collectively, these data indicate that LB-100 results in alternative recognition of introns and exons in human colorectal carcinoma cells, an effect that coincides with potent changes in the phosphorylation status of cancer-critical splicing factors.

### PP2A inhibition-dependent alternative splicing decreases protein expression of resulting transcripts

Alternative splicing impacts the fate of resulting transcripts in various ways, ranging from changed stability, through the formation of premature termination codon (PTC), creation of alternative protein isoform that would differ in its localization, stability or function, or differential translation rates resulting in changed protein levels^23^. To understand the implications of AS changes on gene expression in response to LB-100, we first sought to determine how alternative splicing events downstream of PP2A inhibition alter the levels of resulting proteins. Strikingly, we observed a significant decrease in protein levels of transcripts produced from alternatively spliced mRNAs compared to the background proteins (Figure 2F). Detailed analysis revealed that this was caused primarily by the group of transcripts with differential exon skipping type of ASE and not by the other groups (Figures 2F and S2A-D). Such effect may be mediated by the inclusion of poison exons harboring STOP codons, by the formation of PTC through splicing-in an out-of-the-frame exon, by the formation of alternative 5’ or 3’ Untranslated Regions (UTRs), or by the inclusion of protein domains with putative ubiquitination sites. While the latter two are predicted to change the protein expression through post-transcriptional control, without concurrent changes in mRNA levels, the former two could result in accompanying drops in transcript levels. Therefore, to address the level at which LB-100 stimulation affected protein abundance of AS transcripts, we performed a time-resolved analysis of gene expression by deep sequencing of mRNAs harvested at different time points after PP2A inhibition (Figures 2G and S2E). Critically, we observed that none of the ASE types resulted in decreased abundances of mRNAs undergoing differential splicing events. Rather, some types resulted in paradoxically elevated levels of alternatively spliced mRNAs (Figure 2G and S2E, F). Thus, we concluded that the observed effects of LB-100 on alternative splicing and decreased expression of resulting proteins are unlikely to be mediated by transcriptional control.

Collectively, these observations entail that LB-100 impacts levels of proteins produced on templates of alternatively spliced mRNAs through post-transcriptional regulation, the precise nature of which will be a subject of future studies.

### Targets of splicing-modulating therapies are differentially phosphorylated in response to PP2A inhibition

Phosphorylation of splicing factors was reported previously to be essential for the progression of the splicing reaction^24^. This includes phosphorylation of SF3B1 at the critical threonine residues within the N-terminus of the protein. Recent study determined CDK11 as a kinase responsible for SF3B1 phosphorylation and transition from B to B^act^ stage of the splicing^25^. This transition renders the spliceosome catalytically competent for the first transesterification reaction. Given that SF3B1 is a primary target of splicing-inhibiting drugs proposed as a possible anti-cancer therapy, we speculated that PP2A inhibition affected SF3B1 phosphorylation at the residues important for splicing reactions. Drugs targeting SF3B1, such as pladienolides or the orally available H3B-8800, are proposed to drive alternate functionality of the SF3 complex, resulting in changed splicing patterns in particularly vulnerable cancer cells^18^. Our analysis revealed that LB-100 stimulation affected phosphorylation within the threonine-proline-rich region, with Ser217, Thr227, and Thr326/328 undergoing hypophosphorylation in response to PP2A inhibition (Figure 3A). Notably, Thr328 is one of the residues previously reported to drive the transition to the B^act^ stage^25^, and all observed differential phosphorylation events were grouped in close proximity to the RBM39 protein-protein interaction domain (Figure 3A). Curiously, RBM39 is a target of indisulam, a ubiquitin ligase modulator that marks this protein for degradation and is a subject of ongoing anti-cancer studies^26^. In our results, RBM39 underwent differential phosphorylation at Ser217 within N-terminus and Ser337 within activating domain/ESR proteins interaction region (Figure 3B). These results indicate that primary anti-cancer targets within the spliceosome are differentially phosphorylated in response to LB-100 treatment.

**Figure 3.**
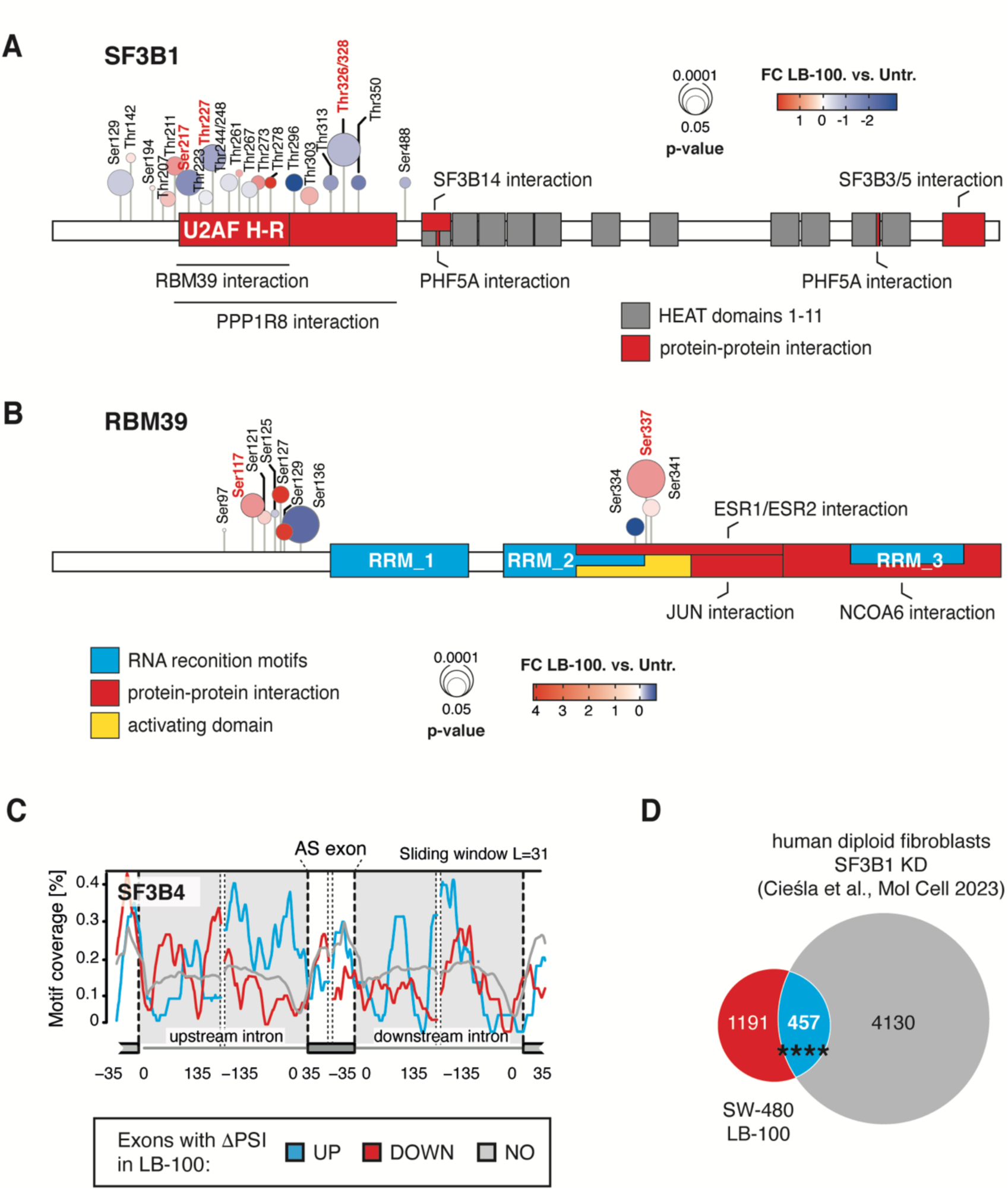
Targets of splicing-targeting drugs are differentially phosphorylated in response to PP2A inhibition. **(A-B)** Location of mis-phosphorylated sites across SF3B1 **(A)** and RBM39 **(B)** proteins. Functionally relevant protein regions according to UniProt are shown. Significantly changing phosphosites in response to LB-100 are highlighted. **(C)** Motif RNA map shows SF3B4 binding in the proximity of excluded (DOWN), included (UP) or not affected (NO) exons in LB-100 as compared to untreated SW-480 colorectal adenocarcinoma cells. **(D)** Venn diagram shows a significant overlap between AS transcripts in LB-100-treated SW-480 cells and SF3B1-depleted MYC-expressing Human Diploid Fibroblasts (HDF) ^20^. ****p<0.0001 (Pearson correlation coefficient). See also Figure S3.

Phosphorylation is predicted to change the functionality and assembly of protein complexes. To determine if the observed phosphorylation changes in spliceosome constituents affected their function, we performed motif enrichment analysis for the binding of these complexes along differentially spliced mRNAs. Indeed, we observed qualitative differences in motif enrichment for SF3B4, a protein acting within the same complex as SF3B1, in between up- and down-spliced mRNAs in LB-100-treated cells (Figure 3C). Moreover, comparative analysis revealed a strong overrepresentation of mRNAs that we previously observed to be alternatively spliced upon knock-down of SF3B1 under oncogenic stress^20^ and transcripts sensitive to LB-100 treatment (Figure 3D). These observations may be widely applicable to other differentially phosphorylated splicing factors, including members of the SRSF family (Figure S3A-C). Collectively, our results propose that phosphorylation changes in splicing factors expose a spliceosome vulnerability in PP2A-inhibited cancer cells.

### LB-100-driven alternative splicing impacts DNA repair regulators and is predicted to result in cancer neoantigens formation

The abnormal splicing evoked by the changed activity of splicing factors could result in increased genotoxic stress, as well as formation of cancer neoantigens recognized by the immune system. Therefore, we set out to determine if LB-100-mediated ASE changes would affect any of these molecular events. Notably, analysis of skipped exon events, which were predicted to change protein expression of resulting transcripts (Figure 2F), revealed a strong enrichment for terms related to the maintenance of genomic integrity, including DNA repair, response to DNA damage checkpoint signaling, or homology-dependent repair (Figure 4A). These observations remained consistent across two different human colorectal adenocarcinoma cell lines and are in agreement both with our findings^20^ and those of others studying SF3B1-inhibited human cells^27^.

**Figure 4.**
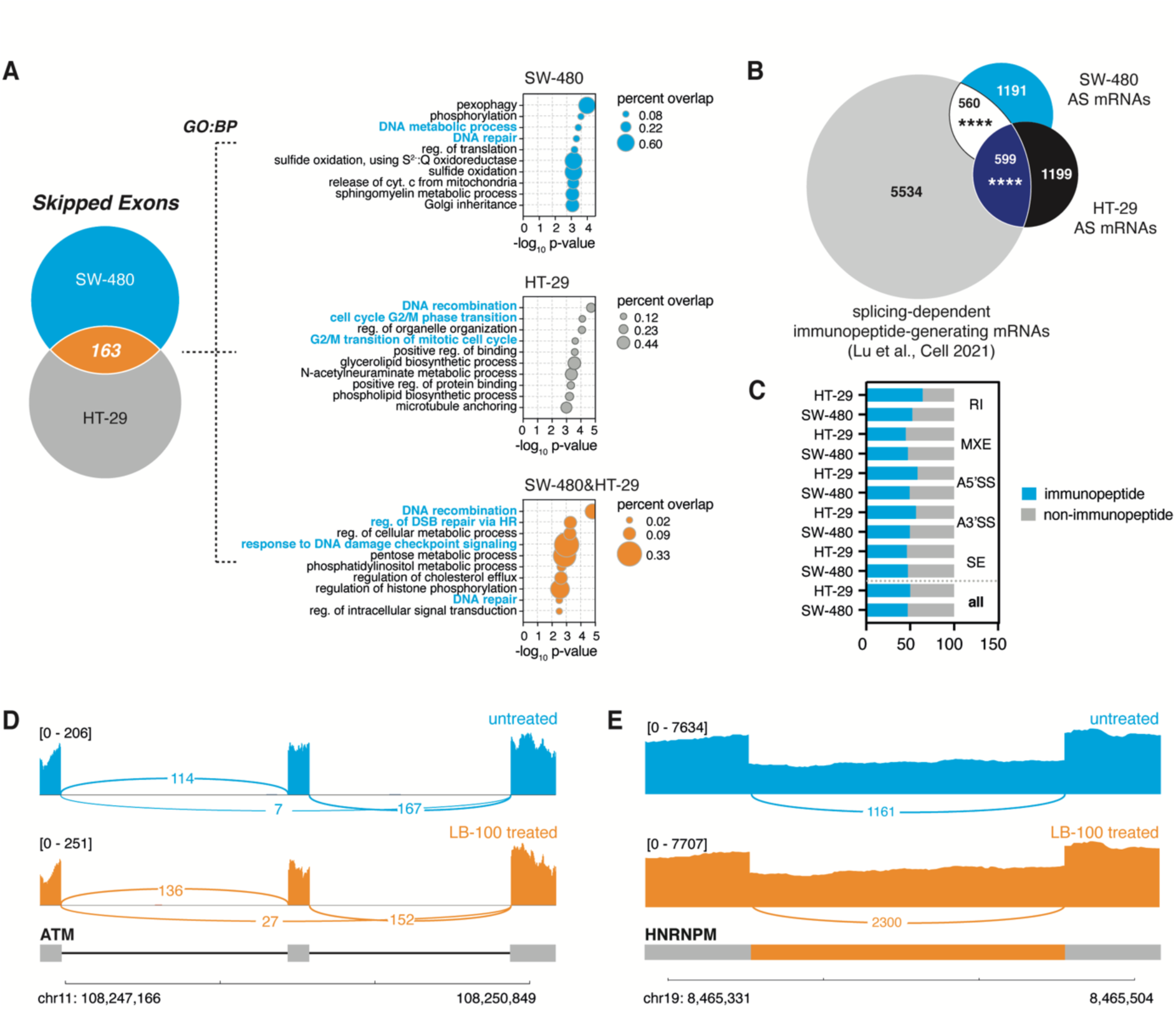
PP2A-sensitive alternative splicing impacts DNA damage related mRNAs to potentially increase neoepitope production. **(A)** Venn diagram shows an overlap between SE in LB-100-treated SW-480 and HT-29 cells (left). Gene Ontology (GO) analysis of SE events in SW-480, HT-29 and an overlap is shown (right). BP – Biological Process. **(B)** Venn diagram shows an overlap between alternatively spliced mRNAs in SW-480 and HT-29 colon adenocarcinomas and previously reported neoantigen-producing mRNAs^14^. ****p<0.0001 (hypergeometric test). **(C)** Bar graph shows equal contribution of all the splicing events toward potential formation of tumor neoantigens reported previously^14^. **(D)** Alternative splicing event within ATM gene and representative Sashimi plots showing differential splicing in untreated and LB-100-treated SW-480 cells. **(E)** Alternative splicing event within HNRNPM gene and representative Sashimi plots showing differential splicing in untreated and LB-100-treated SW-480 cells. See also Figure S4.

Recent evidence describes that splicing inhibition downstream of targeting SF3B1 and RBM39 results in an immune system-dependent extension of lifespan in tumor-bearing mice^14^. Accordingly, we observed that mRNAs that were alternatively spliced in response to LB-100 treatment strongly overlapped with reported immunopeptide-generating ASEs (Figure 4B). Indeed, we predicted that on average, 50% of the alternatively spliced mRNAs from all ASE types could generate cancer neoantigens (Figure 4C), consistent with previous findings^14^. Notably, we observed that alternative splicing impacted on ASE within notable examples of DNA damage response regulators, including apical kinase ATM or PARPBP (Figures 4D and S4A) and previously validated neoantigen-generating mRNA of HNRNPM (Figure 4E). These observations confirm PP2A as a prime anti-cancer target and place LB-100 as a prospective compound for splicing-targeting therapy aiming to enhance the efficacy of genotoxic stressors or immunotherapy approaches.

## DISCUSSION

The current work identifies a major role for PP2A in the regulation of RNA splicing. Our data indicate that inhibition of PP2A with the small molecule inhibitor LB-100 leads to major changes in the phosphorylation of splicing proteins, resulting in aberrant RNA splicing events, such as skipped exons and retained introns. This impact of phosphorylation of splicing factors by PP2A inhibition is supported by other phosphoproteomic analyses^28^. Our findings also indicate that PP2A inhibition induces adverse splicing events in colorectal cancer cells, which overlap with events previously reported to generate neoantigens recognized by cytotoxic T cells. For instance, studies involving the molecular glue indisulam, which causes selective degradation of the splicing factor RBM39^26^, demonstrated that the resulting mis-splicing events create neoantigens, thereby enhancing the efficacy of immune checkpoint blockade^14^. Interestingly, we found that two major phosphorylation sites on RMB39 are affected by LB-100, corroborating the possibility that LB-100, like indisulam, may enhance the efficacy of immune checkpoint blockade. Indeed, multiple studies in a variety of cancer models have demonstrated that LB-100 is synergistic with immune checkpoint blockade^8–10^. Furthermore, patients carrying a mutation in the PP2A scaffold protein PP2R1A have been reported to exhibit the superior response to immune checkpoint blockade therapy^10^. However, it is uncertain at this point whether the observed synergy between LB-100 and immune checkpoint blockade can be attributed to generation of neoantigens as a result of adverse splicing events, as LB-100 also has other immune modulatory effects. For example, it was shown that ablation of PP2A in regulatory T cells (Tregs) inhibits their immunosuppressive activity and causes auto-immunity^29^. Moreover, PP2A inhibition in glioma activates the cGAS-STING pathway, leading to activation of interferon signaling and an increase in CD8+ killer T cell proliferation^30, 31^. The role of PP2A in increased T cell proliferation has also been observed by others^32^. Additional work highlights the potential of altering PP2A activity in conjunction with CDK9 inhibition^33^, with CDK9 being a previously reported modulator of splicing programs^34^. Together, these data provide a strong rationale for testing the combination of LB-100 and immune checkpoint blockade in the clinic. A first trial testing this was started recently (https://classic.clinicaltrials.gov/ct2/show/NCT04560972).

## Acknowledgments

We thank members of the Bernards and Cieśla laboratories for helpful discussion and thoughtful feedback. We thank the Genomics core facility of the Netherlands Cancer Institute for RNA-seq experiments and the Protein facility for carrying out the proteomics experiments. We acknowledge Michał Krzysztoń for critical input on data analysis. This work was supported by an institutional grant of the Dutch Cancer Society and of the Dutch Ministry of Health, Welfare and Sport and by the Oncode Institute (R.B) and by a research grant from Lixte Biotechnology (R.B). M.C. is beneficiary of Sonata Bis (UMO-2022/46/E/NZ3/00141) and OPUS (UMO-2021/43/B/NZ3/01177) grants from National Science Center in Poland. O.B.B. received support of the X-omics Initiative (Project 184.034.019), part of the NWO National Roadmap for Large-Scale Research Infrastructures.

## Contributions

Conceptualization, R.B. and M.C.; Methodology, M.H.D., O.B.B., and V.L..; Investigation, M.H.D., and O.B.B.; Resources, O.B.B., A.V.; Software, A.V. and V.L.; Formal Analysis, M.D.H, V.L., R.B., and C.B.; Data Curation, A.V. and V.L.; Writing - Original Draft, M.D.H., R.B. and M.C.; Writing – Review & Editing, M.D.H., V. L., A. W., R. B. and M.C.; Supervision, R.B. and M.C.; Project Administration, R.B. and M.C.; Funding acquisition, R.B.

## Conflict of interest

Rene Bernards received research funding from Lixte Biotechnology Holdings, the company that manufactures LB-100. Rene Bernards is also a member of the board of directors of Lixte Biotechnology Holdings Ltd. M.H.D. is a shareholder of Lixte Biotechnology. Other Authors have not reported any conflict of interests.

---

**Figure S1.**
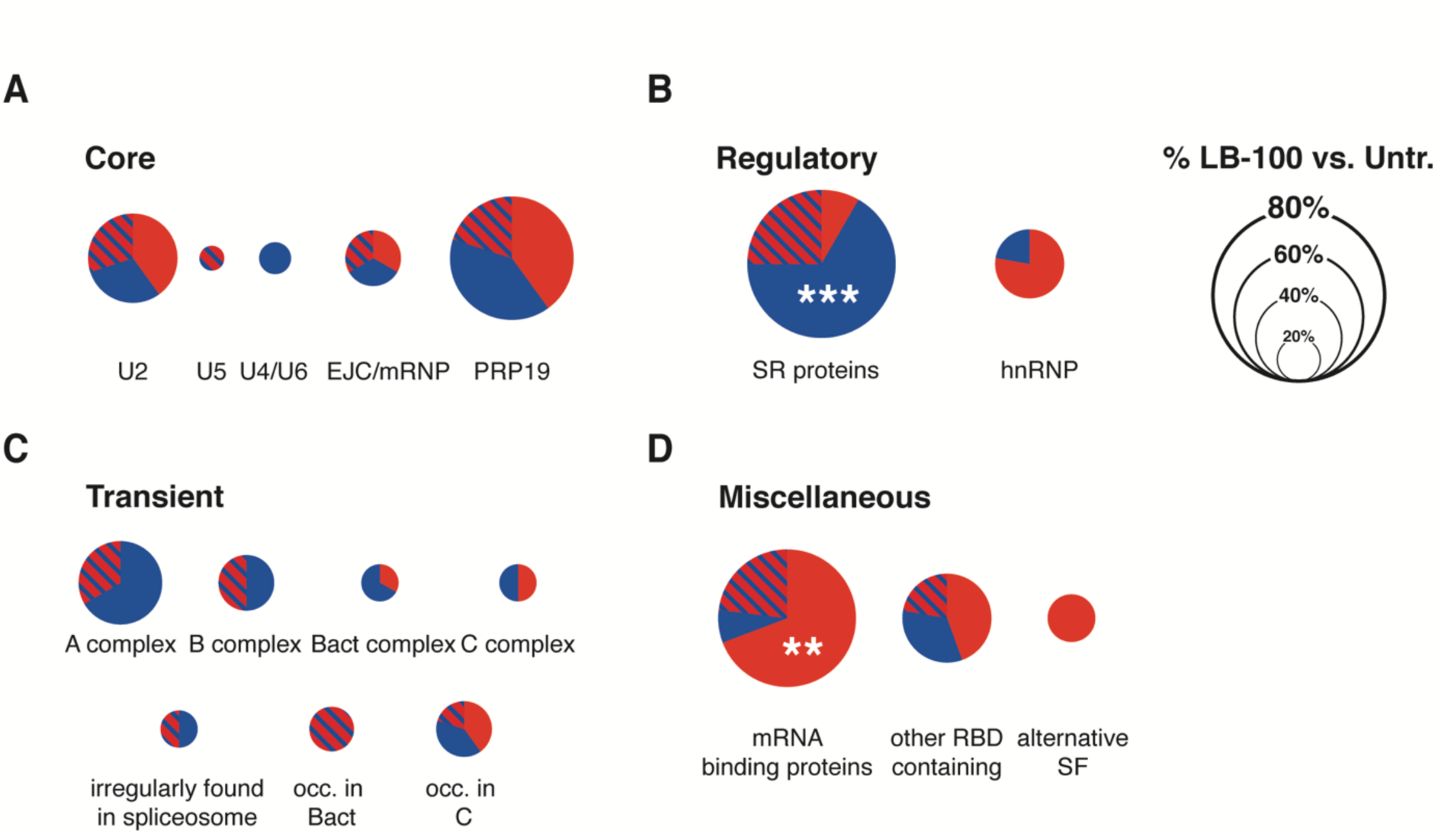
Specific splicing nodes are differentially phosphorylated in response to LB-100 treatment in colorectal adenocarcinoma. Pie charts show distribution of proteins with significantly changed (FDR<0.05 and |Log_2_Fold Change|>0.58) phosphorylation status within different functional groups of splicing factors. ****p<0.0001 (hypergeometric test vs. background). RBD – RNA binding domain; SF – splicing factor. Related to Figure 1

**Figure S2.**
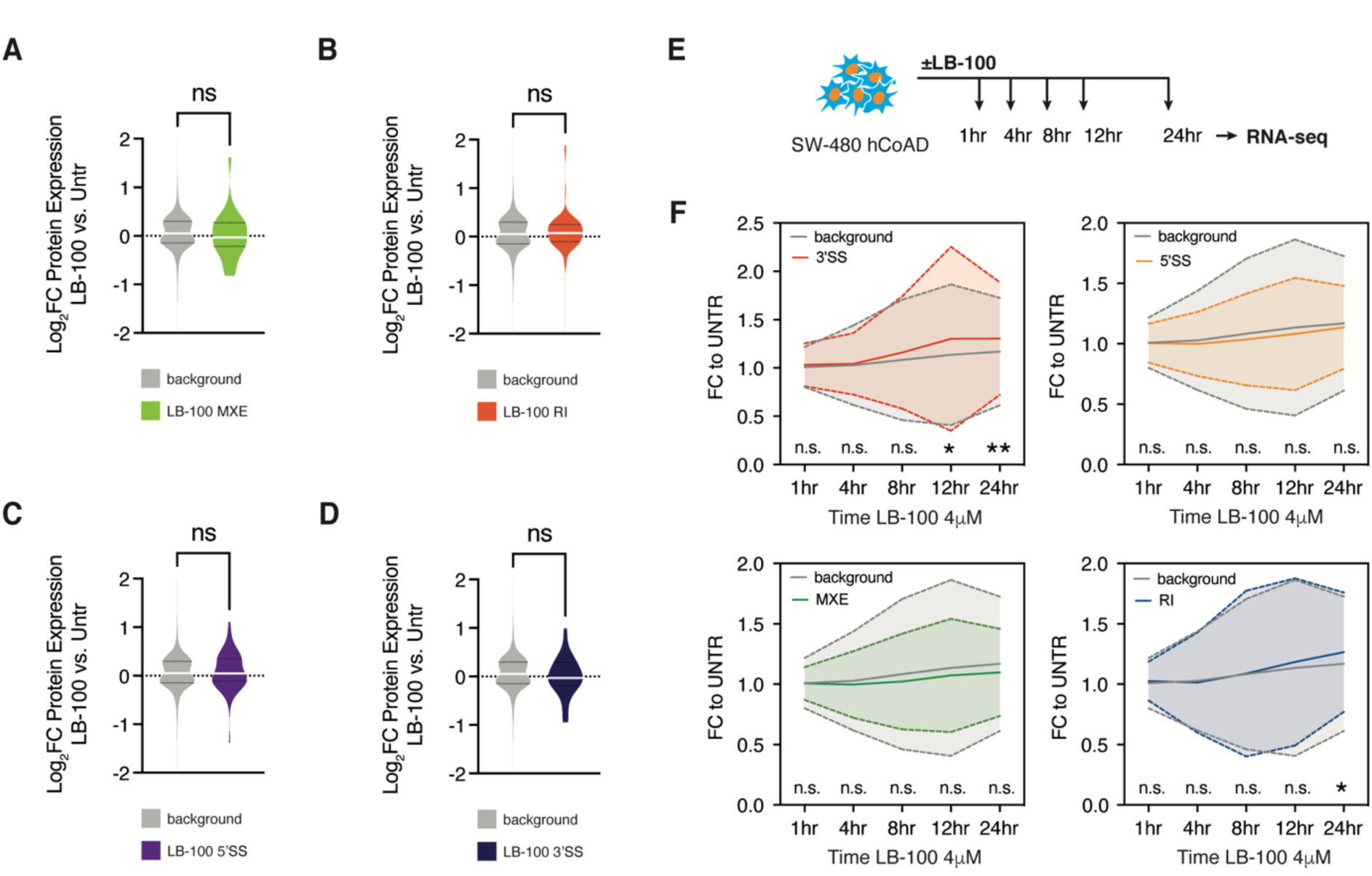
PP2A inhibition-dependent alternative splicing affects protein but not transcript levels of alternatively spliced mRNAs. Violin plots show no effect on levels of proteins resulting from alternatively spliced transcripts in MXE **(A)**, RI **(B)**, alt5’ss **(C)**, and alt3’ss **(D)** in 4 μM LB-100-treated SW-480 cells compared to control. **p<0.01 (t test). **(E)** Schematic illustrates experimental approach to determine time-resolved transcriptomic changes of alternatively spliced mRNAs in response to 4 μM LB-100 stimulation. **(C)** Graphs show time-resolved changes in alternatively spliced transcripts and mRNAs from RI, MXE, alt5’ss and alt3’ss groups in response to LB-100 stimulation. **p<0.01; *p<0.05 (one-way ANOVA). Related to Figure 2

**Figure S3.**
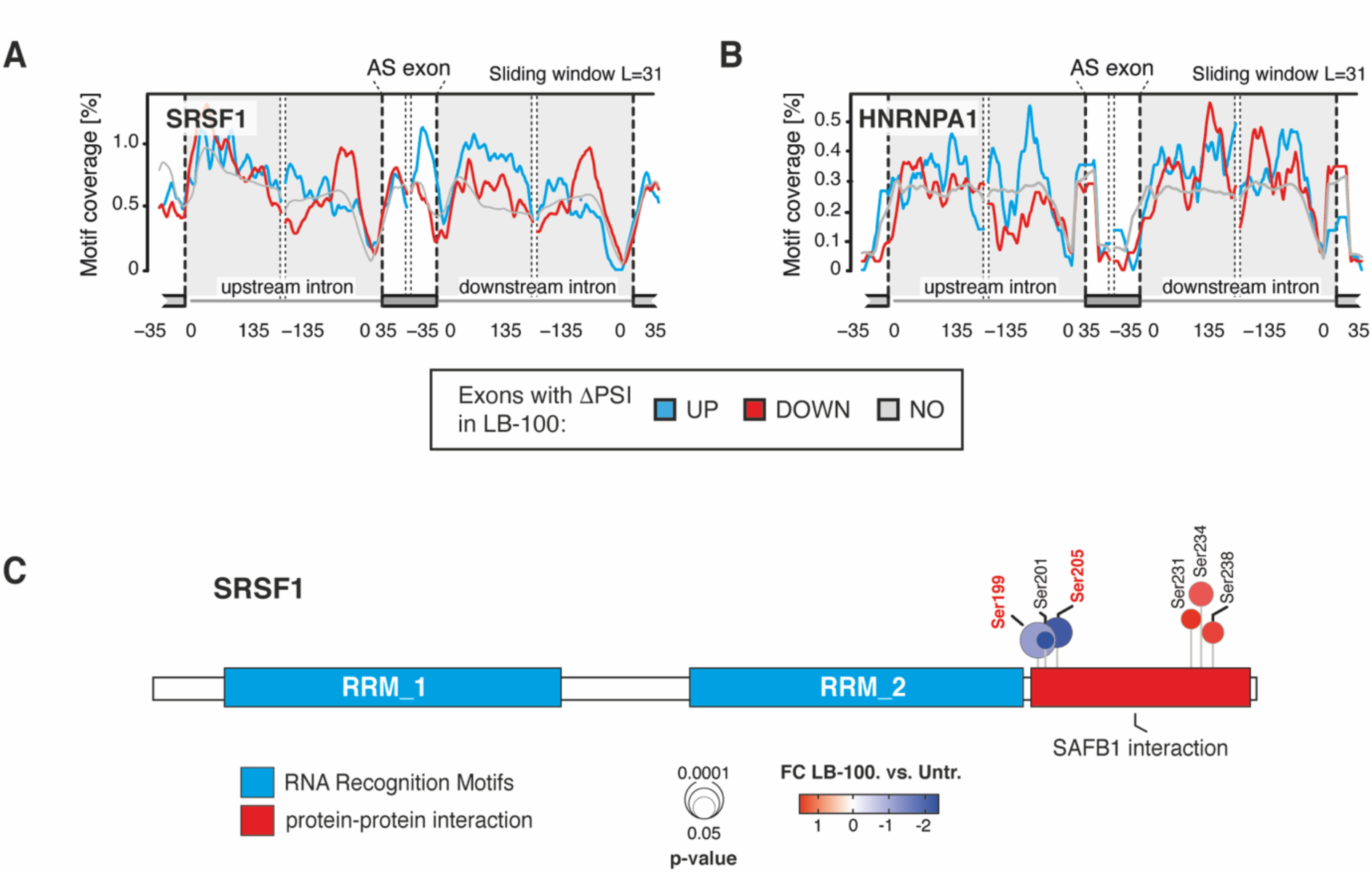
Impact of LB-100-driven phosphorylation changes on function of spliceosome components. **(A-B)** Motif RNA map shows SRSF1 (A) and HNRNPA1 (B) proteins binding in the proximity of excluded (DOWN), included (UP) or not affected (NO) exons in LB-100-treated as compared to untreated SW-480 colorectal adenocarcinoma cells. **(C)** Location of mis-phosphorylated sites across SRSF1 protein. Functionally relevant protein regions according to UniProt are shown. Significantly changing phosphosites in response to LB-100 are highlighted. Related to Figure 3

**Figure S4.**
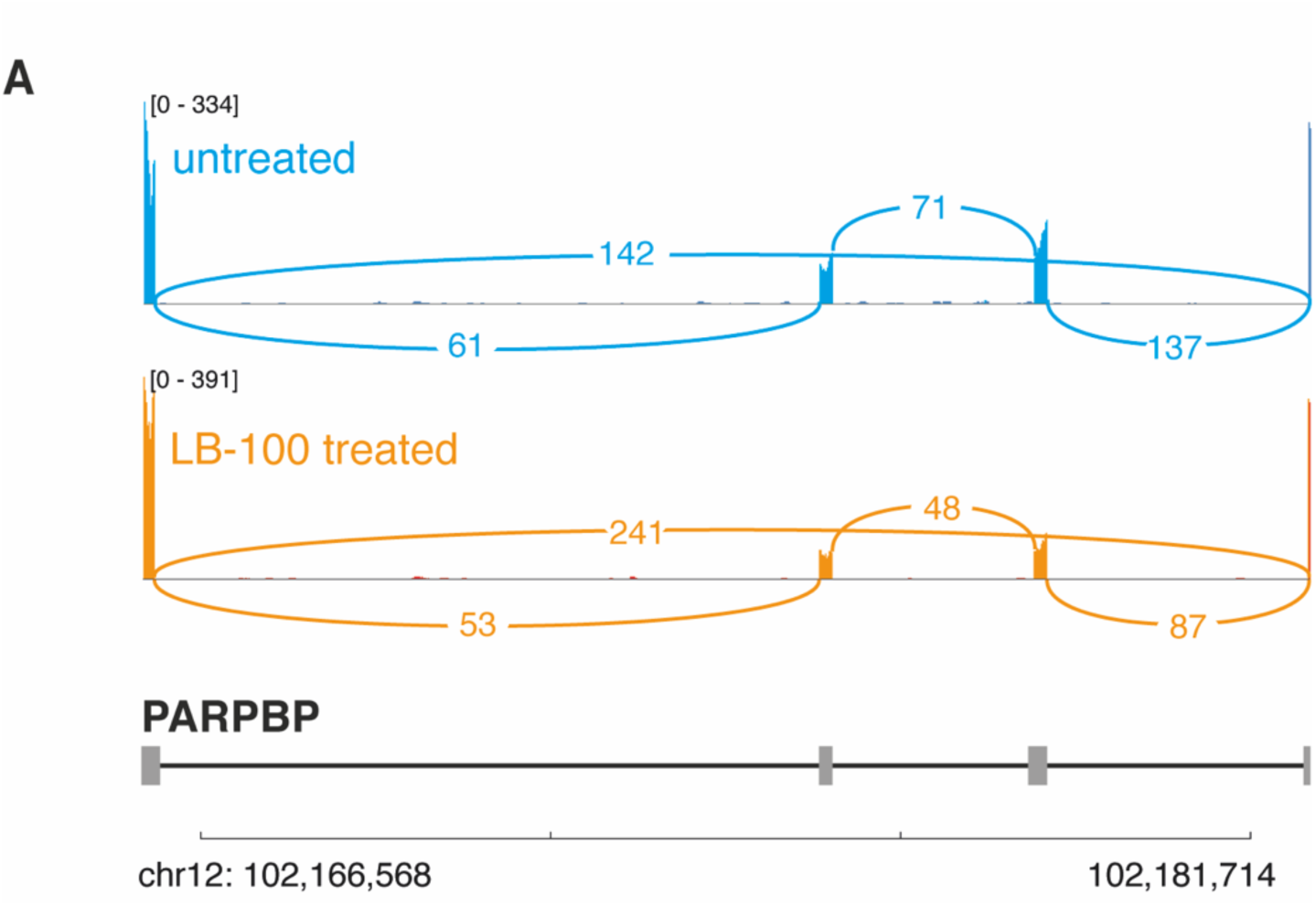
Alternative splicing events in response to LB-100 impact the fate of DNA damage response regulators. **(A)** Alternative splicing event within PARPBP gene and representative Sashimi plots showing differential splicing in untreated and LB-100-treated SW-480 cells. Related to Figure 4

**Table S1** Related to Figures 1 and S1. Proteomic analysis of LB-100 (4μM for 12 hrs) treated SW-480 cells.

**Table S2** Related to Figure 1. Phosphoproteomic analysis of LB-100 (4μM for 12 hrs) treated SW-480 cells.

**Table S3** Related to Figures 2 and S2. Alternative splicing events (ASEs) in LB-100-treated SW-480 and HT-29 colon adenocarcinoma cells determined by RNA-seq. Significant ASEs with |ΔPSI|>0.1 are listed.

**Table S4** Related to Figures 2 and S2. Differential gene expression in response to time-resolved inhibition of PP2A by LB-100 in SW-480 colorectal adenocarcinoma cells.

## EXPERIMENTAL MODEL AND SUBJECT DETAILS

### Lead Contact

Further information and requests for resources should be directed to and will be fulfilled by the Lead Contact, Maciej Cieśla.

### Materials availability

All unique/stable reagents generated in this study are available from the lead contact with a completed Materials Transfer Agreement.

### Data and code availability

RNA-seq data have been deposited to GEO database (https://www.ncbi.nlm.nih.gov/geo/) under the following accession numbers: GSE236625. Proteomics data have been deposited to the ProteomeXchange Consortium via the PRIDE ^36^ partner repository with identifier PXD043658.

## EXPERIMENTAL MODEL AND SUBJECT DETAILS

### Cell culture

Human colorectal adenocarcinoma cells SW-480 and HT-29 were purchased from ATCC and maintained in, respectively, Leibovitz’s L-15 Medium and McCoy’s 5A medium supplemented with 10% fetal bovine serum (FBS) (Thermo Fisher Scientific) and 1% penicillin/streptomycin (Thermo Fisher Scientific). All cells were grown at 37°C with 5% CO_2_ and routinely tested for mycoplasma infection (Universal Mycoplasma Detection Kit, ATCC).

### RNA-sequencing

Transcriptome-wide splicing analysis was performed using SW-480 and HT-29 cell lines. These cells were treated with LB-100 (4 µM) for 12 hrs, with untreated cells serving as a reference control for the analysis. Total RNA from the cells was isolated using the RNeasy Mini Kit (Qiagen), which included an on-column DNase digestion (Qiagen), according to the manufacturer’s instructions. The quality and quantity of isolated RNA were assessed using the RNA 6000 Nano LabChip Bioanalyzer (Agilent Technologies). Sequencing libraries were constructed from total RNA using the TruSeq Stranded mRNA library kit (Illumina). The quality of the resulting mRNA libraries was evaluated using a 2100 Bioanalyzer instrument, following the manufacturer’s protocol specified for the Agilent DNA 7500 kit (Agilent Technologies). After dilution to 10 nM and equimolar pooling into multiplex sequencing pools, paired-end sequencing was performed on the NovaSeq 6000 (Illumina) sequencing instrument.

### Proteomics LC-MS/MSmass spectrometry

Frozen LB-100- or control-treated SW-480 cell pellets were lysed in Guanidine (GuHCl) lysis buffer as described ^37^. Lysate protein concentrations were determined with a Pierce Coomassie (Bradford) Assay Kit (Thermo Scientific) according to the manufacturer’s instructions. Aliquots corresponding to 1100 µg of protein were diluted to 2M GuHCl and digested twice (4h and overnight) with trypsin (Sigma-Aldrich) at 37°C, enzyme/substrate ratio 1:75. Digestion was quenched by the addition of TFA (final concentration 1%), after which the peptides were desalted on a Sep-Pak C18 cartridge (Waters, Massachusetts, USA). From the eluates, 50µg-aliquots were collected for proteome analysis, the remainder being reserved for phosphoproteome analysis. Samples were vacuum dried and stored at -80°C until LC-MS/MS analysis or phosphopeptide enrichment. Phosphopeptides were enriched using the High Select Phosphopeptide Enrichment Kits (Pierce) according to the manufacturer’s instructions, after which eluates were vacuum-dried until LC-MS/MS.

Single-shot LC-MS/MS of proteome samples was performed by nanoLC-MS/MS on an Orbitrap Exploris 480 mass spectrometer (Thermo Scientific) connected to a Proxeon nLC1200 system. Peptides were directly loaded onto the analytical column (ReproSil-Pur 120 C18-AQ, 2.4μm, 75 μm × 500 mm, packed in-house) and eluted in a 90-minutes gradient containing a linear increase from 6% to 30% solvent B (solvent A was 0.1% formic acid/water and solvent B was 0.1% formic acid/80% acetonitrile). The Exploris 480 was run in data-independent acquisition (DIA) mode, with full MS resolution set to 120,000 at m/z 200, MS1 mass range was set from 350-1400, normalized AGC target was 300% and maximum IT was 45ms. DIA was performed on precursors from 400-1000 in 48 windows of 13.5 Da with an overlap of 1 Da. Resolution was set to 30,000 and normalized CE was 27.

For single-shot LC-MS/MS of phosphoproteome samples, a 135-min gradient containing a 114-min linear increase from 7 to 30% solvent B was used. The Exploris 480 was run in data-dependent acquisition (DDA) mode with full MS resolution set to 120,000 at m/z 200 and a cycle time of 2 sec. MS1 mass range was set from 375-1500, normalized AGC target was 300% and maximum IT mode was set to ‘Auto’. Exclusion duration was set to 30 sec; MS2 spectra were acquired at 30,000 resolution with data dependent mode set to ‘cycle time’. Precursors were HCD-fragmented with a normalized collision energy of 27 when their charge states were 2-6; MS2 isolation window was 1.2 m/z, the normalized AGC target was set to “standard” and the maximum injection time mode was set to “auto”.

### RNA-seq and splicing analysis

Paired-end sequencing was performed using 101 cycles for Read 1, 19 cycles for Read i7, 10 cycles for Read i5 and 101 cycles for Read 2, using the NovaSeq 6000 SP Reagent Kit v1.5 (200 cycles) (20040719, Illumina). Sequencing data was demultiplexed into FastQ files using BCLConvert version 3.9.3 (Illumina). The paired-end reads were trimmed for adapter sequences using SeqPurge version 2019_09 ^38^ and aligned to the human hg38 reference genome using HiSat2 ^39^ version 2.1.0 with the pre-built grch38_snp_tran reference. Gene counts were generated using HTseq-count ^40^ and Homo_sapiens.GRCh38.102.gtf. Splicing events were determined using rMATS version 4.1.2 ^22^ using the Ensembl release 100 transcript GTF ^41^. An ASE was considered significant based on cut-off for FDR<0.05 and |ΔPSI|>0.1 for the individual condition supporting existence of transcript with ASE. Analysis of sequence features associated with identified ASEs was performed with MATT version 1.2.1 ^42^, comparing 410 LB-100-induced (UP) and 641 LB-100-repressed (DOWN) skipped exons or 87 (UP) and 105 (DOWN) retained introns to the background (NO). The enrichment for RNA binding proteins sites within LB-100-regulated exons was calculated using the CISBP-RNA binding motif RNA-maps ^43^ using default settings. The following groups were used for comparison: exons with increased (UP) and decreased (DOWN) inclusion upon LB-100 treatment of SW-480 cells, and reference group (NO; FDR>0.05 and |ΔPSI|>0.1). Gene Ontology (GO) analysis was performed using Enrichr.

### Proteomics data analysis

Proteome data were analyzed with DIA-NN (version 1.8) ^44^ without a spectral library and with “Deep learning” option enabled. The Swissprot Human database (20,395 entries, release 2021_04) was added for the library-free search; the quantification strategy was set to ‘Robust LC (high accuracy)’ and the MBR option was enabled. All other settings were kept at the default values. The protein groups report from DIA-NN was used for downstream analysis in Perseus (version: 1.6.15.0) ^45^. Values were Log2-transformed, after which proteins were filtered for at least 75% valid value presence in at least one sample group. Missing values were replaced by imputation based a normal distribution using a width of 0.3 and a minimal downshift of 2.4. Differential protein abundances were determined using a student’s t-test (minimal threshold: FDR: 5% and S0: 0.35). Phosphoproteome data were analyzed using MaxQuant (version 2.0.3.0) ^46^ using standard settings for label-free quantitation (LFQ). MS/MS data were searched against the same sequence database as mentioned above, complemented with a list of common contaminants and concatenated with the reversed version of all sequences. The maximum allowed mass tolerance was 4.5ppm in the main search and 0.5 Da for fragment ion masses. False discovery rates for peptide and protein identification were set to 1%. Trypsin/P was chosen as cleavage specificity allowing two missed cleavages. Carbamidomethylation was set as a fixed modification, with methionine oxidation and Phospho(STY) as variable modifications. Phosphosite LFQ intensities were were extracted from the Phospho(STY)sites.txt file and processed in Perseus as described above and filtered for reverse sequences and potential contaminants, as well as for site probability ≥0.75. Additionally, sites were filtered for at least 100% valid value presence in at least one sample group. Values were normalized by median substraction and missing values were replaced by imputation based on a normal distribution (width: 0.3 and downshift: 1.8). Phosphosites with differential abundance were determined using a Student t-test (minimal threshold: FDR: 5% and S0: 0.1). All mass spectrometry proteomic data generated in this study have been deposited to the ProteomeXchange Consortium via the PRIDE ^36^ partner repository.

### Quantification and statistical analysis

Data is presented as mean, ± SD or SEM, unless otherwise stated. Indicated number of independent biological replicates have been performed for each experiment. Statistical tests used and specific p-values are indicated in the figure legends.

